# Marker effect p-values for single-step SNP-BLUP genomic models

**DOI:** 10.1101/2025.10.30.685527

**Authors:** Damilola Adekale, Zengting Liu, Hatem Alkhoder, Dierck Segelke, Georg Thaller, Jens Tetens

**Affiliations:** Functional Breeding – Genetik und züchterische Verbesserung funktionaler Merkmale, Göttingen, Germany; Biometrie, Vereinigte Informationssysteme Tierhaltung w.V., Verden, Germany; Institut für Tierzucht und Haustiergenetik, Christian-Albrechts-Universität zu Kiel

## Abstract

**Background:** Single-step Single Nucleotide Polymorphism best linear unbiased prediction (ssSNPBLUP) is a comprehensive method for obtaining genomically enhanced breeding values for animals and SNP effects in a single evaluation. The ssSNPBLUP model integrates phenotypic, pedigree, and genomic data for genomic evaluations. However, there has been no framework for estimating the reliability and p-values of the SNP effects obtained from a ssSNPBLUP genomic model. This study investigates the reliability and significance of the SNP effects estimated using a ssSNPBLUP framework in German Limousin (LIM) and Holstein (HOL) cattle populations.

**Methods:** This study introduces a novel approach for calculating p-values within the ssSNPBLUP framework and compares it to a conventional single-marker regression GWAS approach. SNP reliabilities were computed using prediction error variances of SNP effect estimates, enabling the identification of statistically significant SNP markers. LIM data included weaning weight (200-DW) evaluated with a maternal effect BLUP model, while HOL data comprised production traits (milk yield, protein yield, fat yield, and somatic cell score) analysed via a random regression test-day model.

**Results:** The results reveal significant SNP effects in both LIM and HOL evaluations, with notable differences attributed to the size of the reference populations. Average SNP reliabilities were higher in HOL (Mean SNP reliability: 0.42) compared to LIM (Mean SNP reliability: 0.02), underscoring the critical role of the size of the reference population in determining the accuracy and reliability of SNP effects obtained from genomic evaluations.

**Conclusions:** The calculation of p-values from the ssSNPBLUP framework offers an efficient approach to identify quantitative trait loci (QTL) that significantly influences traits in populations. Our approach provides a framework that could be implemented in large and complex datasets such as those used in many national routine evaluations, where only a proportion of the animals are genotyped.

## Background

Understanding and predicting genetic risk factors for essential traits requires understanding the specific loci underlying a phenotype and a trait’s genetic architecture. Conventional approaches to detect the genomic regions associated with phenotypes involve single marker regression [1,2]. In traits with simple genetic architecture, single marker regression approaches such as Genome-wide association Studies (GWAS) are an efficient way to discover QTLs associated with phenotypes. Conventional single marker regression GWAS approaches sequentially fit all SNPs as an effect in a mixed model equation that includes the pedigree or genomic relationship matrix to adjust for population structure and relationship. However, many traits often possess more complex architectures that present difficulties for GWAS. A common issue with conventional GWAS approaches is the missing heritability where identified variants only explain a small portion of phenotypic variance. A consensus in the literature is that complex traits are caused by a large number of variants, the effects of which are too small to pass a stringent genome-wide significance level [3]. Further complicating the applicability of conventional GWAS approaches to livestock, conventional GWAS approaches usually require all individuals to have a phenotype and genotype record which is unfeasible in many livestock populations due to cost and biological limits. Deregressed proofs have been adopted for GWAS to detect QTLs in cases such as sex-limited traits [4,5]. However, depending on the approach, deregressed proofs involve significant approximation and projection of phenotypes from relatives. The significance of genetic markers on a trait depends on the linkage disequilibrium between the SNP and the QTLs influencing the trait of interest. Considering that single marker regression GWAS approaches simultaneously test for the significance of markers, they do not account for linkage disequilibrium between the SNP markers in proximity to one another. Although single marker regression approaches provide ease of significance testing, it is likely to result in reduced fit to the data compared with methods where all SNPs are jointly considered [4]. Furthermore, the sequential testing of SNPs creates a multiple-testing problem, where the chance of observing false positives increases [6]. To adjust for multiple tests, GWAS approaches have adopted threshold-correcting approaches such as Bonferroni correction [7].

Single-step approaches enable the simultaneous estimation of breeding values of genotyped and ungenotyped animals in a single evaluation. The ssSNPBLUP approaches enable the joint estimation of genetic merit and marker effects while accounting for genetic structure within the population. The estimation of SNP effects in the ssGBLUP approach involves back solving of animal effects to obtain SNP effects [8,9]. Considering the infinitesimal allele model appropriate approaches to detect genomic regions associated with traits should involve all SNPs in a single mixed model equation. Single-step genomic evaluation continues to be adopted in routine genetic evaluation of livestock species [10–12]. The ssSNPBLUP approach enable the simultaneous estimation of breeding values and SNP marker effects from phenotypic information of genotyped and ungenotyped animals in a single evaluation while accounting for genetic structure within the population. The estimation of SNP effects in the ssGBLUP approach involves back solving of animal effects to obtain SNP effects [8,9]. Considering the infinitesimal allele model, appropriate approaches to detect genomic regions associated with traits should involve all SNPs in a single mixed model equation. Single-step approaches overcome the issues faced by conventional single marker GWAS approaches by jointly estimating marker effects and adjusting genetic and population structure in a mixed model equation. However, considering the joint estimation of SNP effects, single-step frameworks lack an approach to compute reliability of SNP effects. A common issue faced is the inability to define thresholds of significance for SNP markers associated with the trait. Several studies have adopted the approach of using explained variance or presenting SNP effects in genetic standard deviations to identify genomic regions affecting the trait of interest [9,11,13]. The estimates of marker effects or explained variances correctly consider the weight of the estimated marker effects but do not always consider the uncertainty in effect estimates. By back-solving breeding values and the prediction error (co)variances, [9] introduced an approach to defining the threshold of significance for the single step GBLUP framework [9,14]. However, to our knowledge, there are no approaches to estimate the reliability of SNP effect estimates for the ssSNPBLUP framework.

This work presents an approach to estimating the uncertainty of SNP effects obtained from a single step SNPBLUP genomic evaluation. We also seek to present significance thresholds for the SNP effects estimates obtained from a single-step SNPBLUP genomic evaluation. We apply this approach to SNP effects from a multiple-trait evaluation of the Limousin German national beef cattle and the milk production traits in the German dairy cattle genomic evaluation. Using the two datasets, we compare the effect of reference population size on the significance of SNPs. We show the method’s applicability to small and large datasets comparable to those used in many national evaluations. Furthermore, using the beef cattle dataset, this study aimed to compare the informative regions identified in a conventional single marker genome-wide association study (GWAS) to regions identified in the single-step SNPBLUP genomic evaluation.

## Method

A single-step SNP BLUP model (ssSNPBLUP) [15] is assumed for computing reliability values of the SNP marker effect estimates. The ssSNPBLUP model is applied to phenotypic records of all cattle for a single-step genomic evaluation:

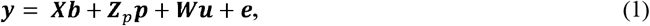

where **y** is an *n*_*T*_ × 1 vector phenotypic records of the cattle, *n*_*T*_ = number of phenotype records; **b** is a *n*_*F*_ × 1 vector of all fixed effects, *n*_*F*_ = number of all fixed effects; **p** is an *n*_*p*_ × 1 vector of non-genetic random effects (e.g. permanent environmental effects of the cows), *n*_*p*_ × 1 = total number of **p** effects; **u** is an *n*_*A*_ × 1 vector of additive genetic effects, *n*_*A*_ = total number of animals (non-genotyped animals denoted as group 1 and genotyped animals as group 2); ***X*** of order *n*_*T*_ × *n*_*F*_, **Z**_***p***_ of order *n*_*T*_ × *n*_*p*_ and of order *n*_*T*_ × *n*_*A*_, are incidence matrices of effects **b, p** and **u**, respectively. As usual, we assume that 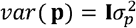 with 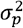 representing variance of the non-genetic random effects. Furthermore, we assume 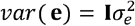 with 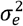 being random error variance.

A residual polygenic effect (RPG) is assumed as in the ssSNPBLUP model [15]. The additive genetic effects of *n* genotyped animals of group 2 can be then separated into two terms:

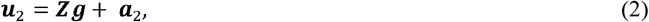

where ***g*** = *m* × 1 vector of additive genetic effects of *m* fitted SNP markers, ***a***_*2*_ = *n* × 1, vector of RPG effects of the genotyped animals, and **Z** is a design matrix of order *n* × *m* containing regression coefficients on genotypes of the reference animals at all the m SNP markers: 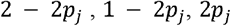 for genotype AA, AB or BB of the j-th marker [16], where *p*_*j*_ represents the allele frequency of the j-th SNP marker. RPG effects of the genotyped animals of group 2 have a normal distribution:

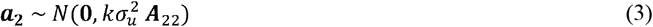

Where **A**_22_ represents pedigree relationship matrix among the genotyped animals. Residual polygenic variance parameter *k*, representing the proportion of additive genetic variance not explained by the *m* SNP marker, may take any value between 0 and 1, but excluding two boundary values *k*=0 and *k*=1. We further assume that additive genetic effects of the SNP markers of the ssSNPBLUP model (Eq. (1)) follow a normal distribution:

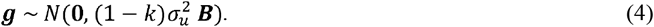

Under the assumption of uncorrelated SNP effects, **B** is a diagonal matrix of order of *m* x *m*. Under the ssSNPBLUP model (Eq. (1)), all the *m* SNP markers explain equal additive genetic variance, thus

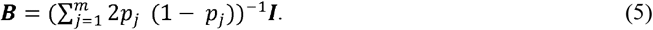

Computing reliabilities of the SNP effect estimates, 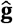, needs their prediction error (co)variances (PEV), which would require inverse of the coefficient matrix of the left-hand-sides (LHS) of mixed model equations (MME) of the ssSNPBLUP model Eq. (1). For a full single-step genomic evaluation like for German Holstein population [17] or for the German beef cattle breeds [11], inverting the full coefficient matrix of the MME is infeasible, because the dimension of the MME system exceeded hundreds of millions for most dairy traits in German Holstein breed. As the number of genotyped animals increases over time rapidly, obtaining the inversion of genomic relationship matrix of ssGBLUP model has also turned out to be impossible. Therefore, an alternative way to approximate the reliabilities of the SNP effect estimates was implemented by applying Interbull’s methods for calculating effective daughter contributions (EDC) of bulls with daughters and effective records contribution (ERC) of cows with phenotypic records. The fixed effects **b** and the non-genetic random effects **p** of Eq. (1) were absorbed stepwise into the additive genetic effects of the cows **u** to obtain ERC of the cows. For bulls with daughters having phenotypic data, ERC of their daughters were further absorbed in the bulls with adjustment for the contribution of dams of the daughters to get EDC of the bulls.

Having completed all the absorption steps above, PEV of the SNP effect estimates of the ssSNPBLUP model (Eq. (1)) can be obtained by inverting LHS coefficient matrix of the following MME:

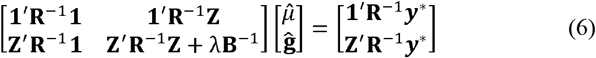

where ***y***\* denote pseudo-phenotypic values for all the genotyped animals with own phenotypic data which are free of all the other effects of model (Eq. (1)) but the additive genetic effects; the pseudo-phenotypic values ***y***\* are not relevant for the computation of reliabilities of the SNP effect estimates 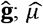 represents a general mean for the sub-population of the genotyped and phenotyped animals; **1** is a vector of 1s with a dimension equal to the number of genotyped animals with phenotypic data; 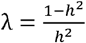 with *h*^2^ being heritability of the evaluated trait; **R**^−1^ is a diagonal matrix **R**^−1^ = *diag* (***φ***), where **φ** represents EDC of genotyped bulls with daughters and/or ERC of genotyped cows with own phenotypic records.

When the genotyped cows and their genotyped sires are both present in the sub-population, EDC of their sires must be adjusted for the contribution of the genotyped cows to avoid double counting of ERC of the genotyped cows [18]. The method implemented in this paper is similar to steps 7 and 8 in [9].

Following [19], we denote the coefficient matrix of MME Eq. (6) by **C** and its inverse matrix element as

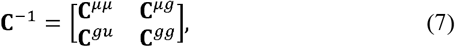

where **C**^*gg*^ is the submatrix associated with the SNP effect estimates. The reliability of marker effect estimate of the *j*-th SNP is calculated as

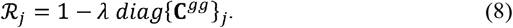

The standard error for marker effect estimate of the *j*-th SNP (*SE*_*j*_) is calculated as

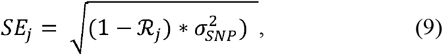

where 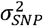 represents average additive genetic variance of the SNP markers:

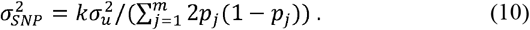

To test the significance of the marker effect, estimate of the *j*-th SNP under the Normal distribution, a so-called Z score is calculated as

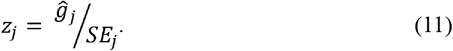

Given the cumulative distribution function of the standard Normal distribution, p-value for effect estimate of the marker *j* is calculated as usual

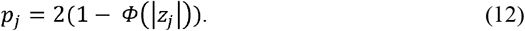

Finally, significant SNPs are identified based on a previously defined significance threshold.

### Conventional GWAS

Using the beef cattle dataset, a conventional pedigree-based evaluation was carried out to estimate the EBVs for the animals. The EBVs are deregressed using the matrix deregression approach [20,21] to calculate the deregressed proofs. Conventional single marker regression GWAS approaches were estimated using GCTA [22].

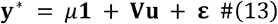

where ***y***\* is an n x 1 vector of 200-DW deregressed proofs, **1** is a vector of ones, *μ* is the general mean, and **u** is a vector of SNP effects with 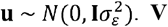 is a standardized genotype matrix with the *ijth* element 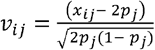, where *x*_*ij*_ is the number of copies of the reference allele for the *jth* SNP of the *ith* individual and *p*_*j*_ is allele frequency of the *j*-th SNP marker [22]. For the p-values in the genomic and single-marker regression approaches, a base rejection threshold of 0.05 and a Bonferroni-adjusted threshold for (0.05/45,613), which equals 5.9 on the −log_10_ scale, were adopted. Due to the large reference population size, a conventional single marker regression GWAS could not be carried out for the HOL dataset.

### Dataset and model

#### Dataset

The German national Limousin (LIM) and Holstein (HOL) datasets for routine breeding value estimation were used in this study. We analysed the SNP effects estimated from the German single-step LIM genomic evaluation [11]. The LIM dataset was obtained from the routine beef cattle evaluation of December 2023. Weaning weight (200-DW) data was considered in this study. The statistical model for the routine genetic evaluation of production traits in German beef cattle is a standard BLUP multi-trait maternal effect animal model [23]. Fixed effects of season of birth, sex, birth type (single or twin), month of birth, and parity nested within the age class of the dam. GEBVs and SNP effects for all traits are simultaneously estimated. We utilised the 200-DW trait in this study. The SNP effects for the HOL dataset were evaluated using a random regression test day model [24]. The model is described in detail by [17]. The p-values of the SNPs in the ssSNPBLUP model were calculated using the abovementioned method. The traits, breed, heritability, genetic variance, and number of reference animals are presented in Table 1.

**Table 1:**
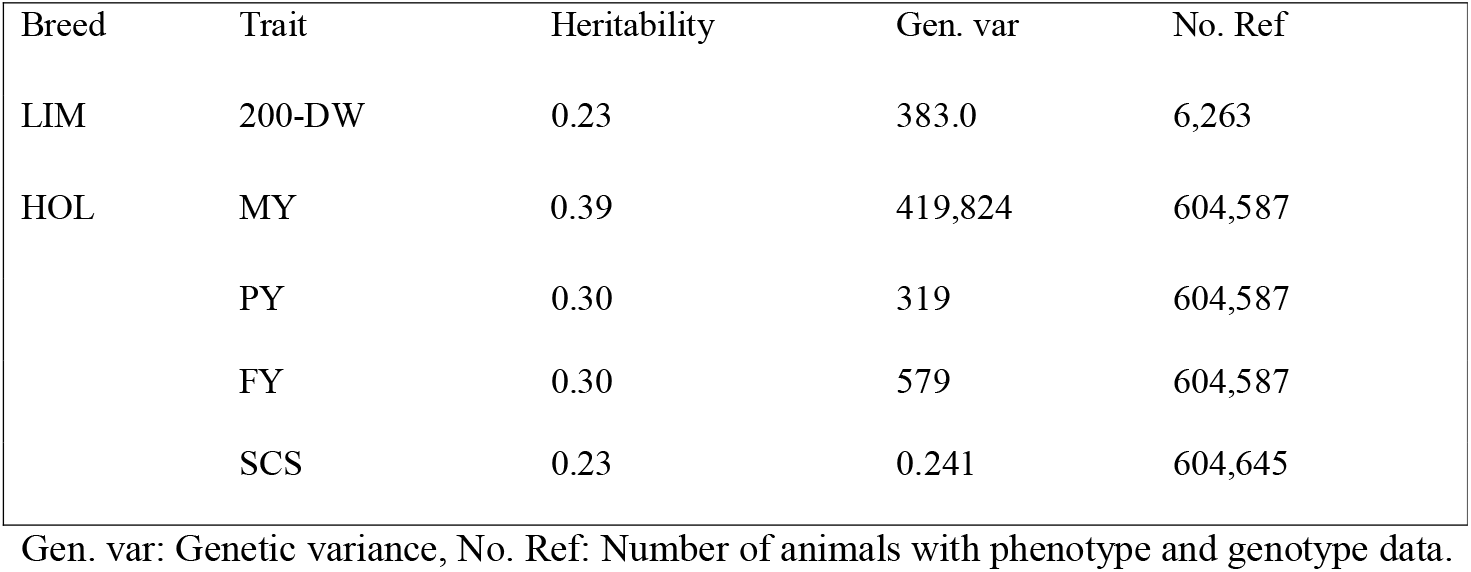
Number of reference animals defined as the number of animals with genotype and phenotype records.

| Breed | Trait | Heritability | Gen. var | No. Ref |
| --- | --- | --- | --- | --- |
| LIM | 200-DW | 0.23 | 383.0 | 6,263 |
| HOL | MY | 0.39 | 419,824 | 604,587 |
|  | PY | 0.30 | 319 | 604,587 |
|  | FY | 0.30 | 579 | 604,587 |
|  | SCS | 0.23 | 0.241 | 604,645 |

Gen. var: Genetic variance, No. Ref: Number of animals with phenotype and genotype data.

The genotypic dataset is based on the Illumina V2 chip and the UMD 3.1 reference genome. The genotypes were imputed to a common panel of 45,613 markers across the 31 chromosomes of cattle. The total number of SNP markers on the autosomes and the sex chromosomes is presented in [17].

The ssSNPBLUP model was implemented in the Mix99 suite of software [25], the single marker regression GWAS approach was implemented in GCTA [22].

## Results

Figure 1 shows the Manhattan plot for the SNP effects (in genetic SD) and associated p-values obtained from the single step genomic evaluation of the LIM evaluation. Figure 2 and Fig. 3 show the Manhattan plot for the SNP effects (in genetic SD) associated p-values for the production traits in HOL, respectively. For the LIM evaluation, the Manhattan plot SNP effects indicate identifying regions of interest on chromosome 6 for DA and LIM. Several studies [26–29] have reported Quantitative Trait Loci (QTL) between 37 and 56 cM of BTA6, affecting body weight, growth, and carcass traits in beef cattle populations. Using the method of calculating p-values described in this paper, Fig. 2 indicates a region of interest for the 200-DW in the LIM evaluation. The p-values obtained from a single marker regression GWAS of the LIM population is presented in Fig. 4. Figure 4 shows SNPs on chromosome 6 that meet the significance threshold in the LIM population. Although, the significant markers in this region have been confirmed in other studies, they failed to meet either the Bonferroni adjusted threshold (0.05/45613) or the base significance threshold (p < 0.05) using the genomic model described in this paper.

**Figure 1:**
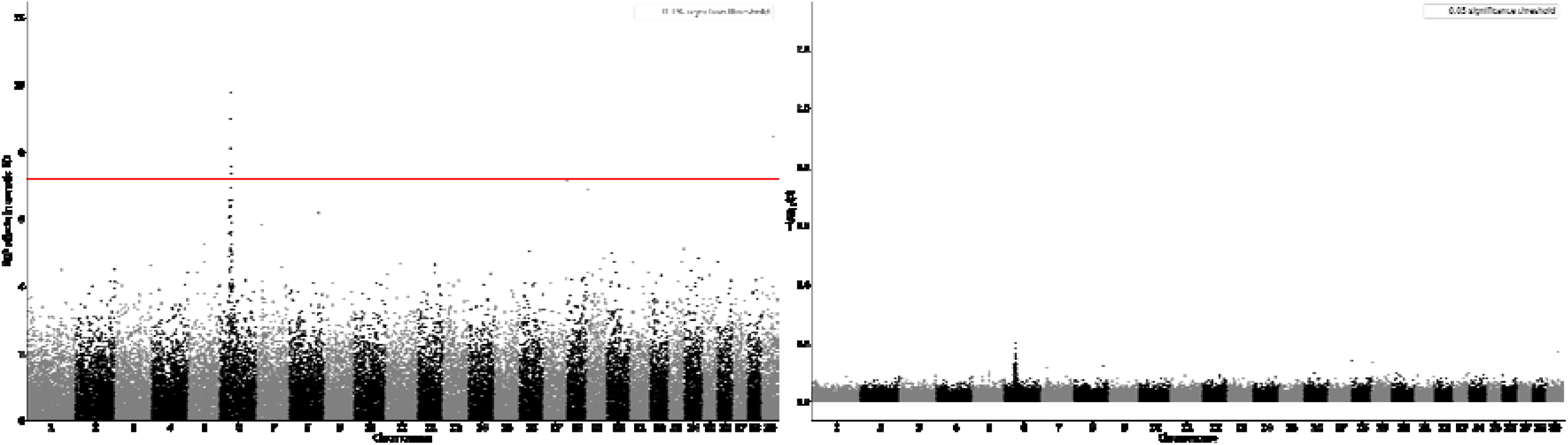
Manhattan plot of SNP effects in genetic SD and p-values for 200-DW in LIM population. **Legend:** SNP effects: SNP effects from genomic model. p-values:

**Figure 2:**
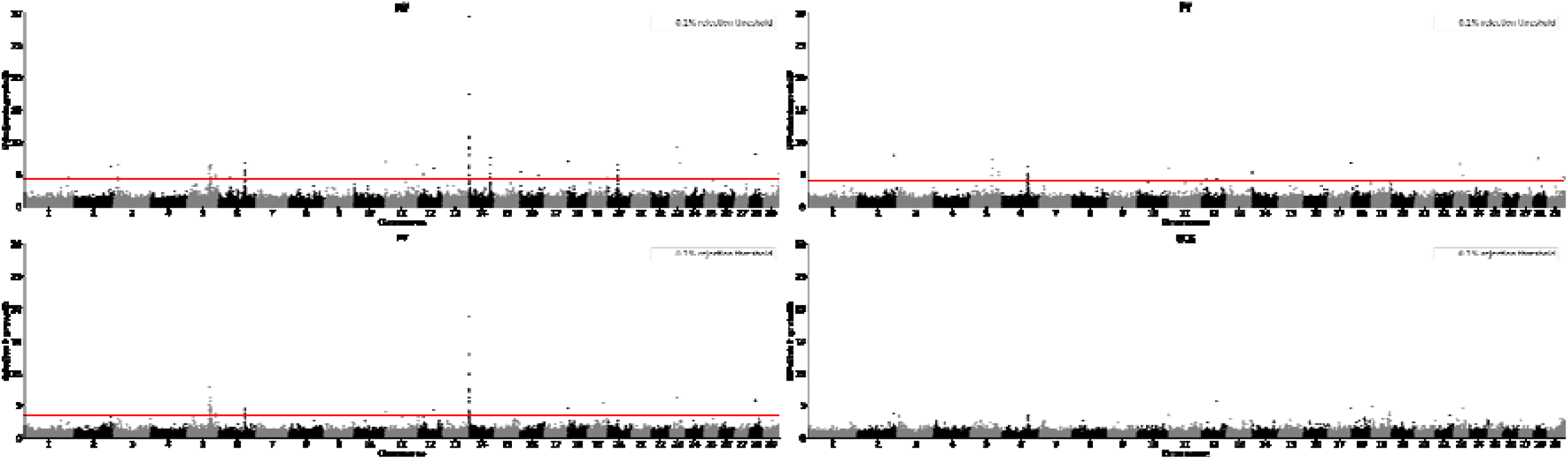
Manhattan plot (SNP effects in genetic SD) from dairy genomic evaluation. **Legend:** MY: milk yield, PY: protein yield, FY: fat yield, SCS: Somatic cell score.

**Figure 3:**
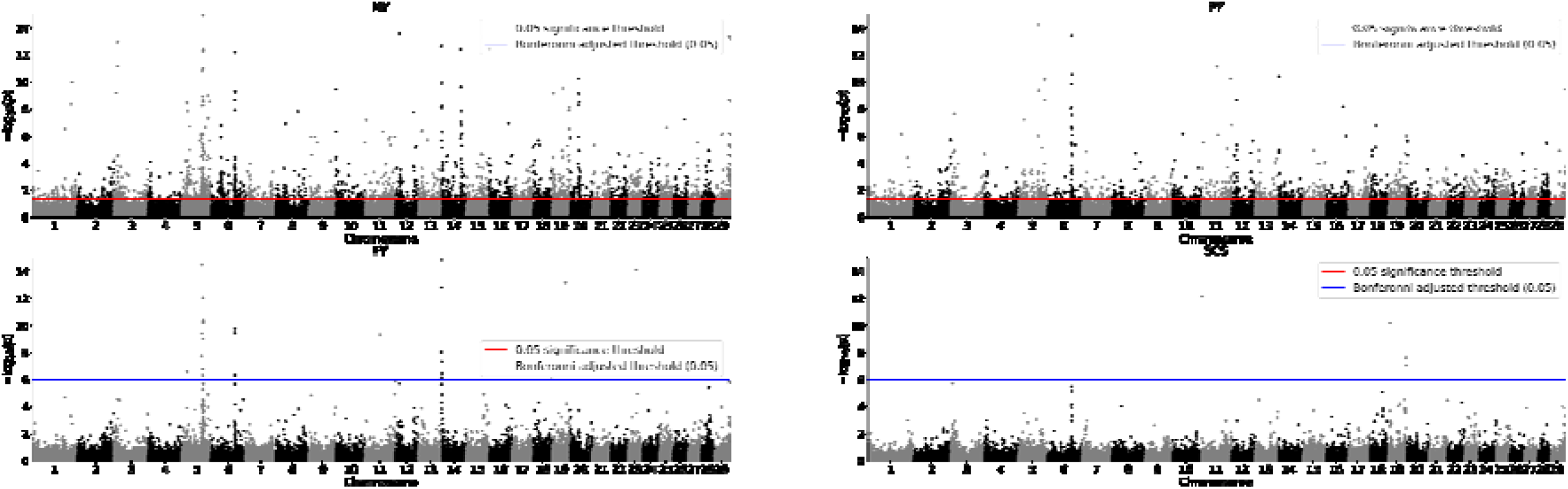
Manhattan plot (p-values) of SNP effects from dairy genomic evaluation. **Legend:** MY: milk yield, PY: protein yield, FY: fat yield, SCS: Somatic cell score.

**Figure 4:**
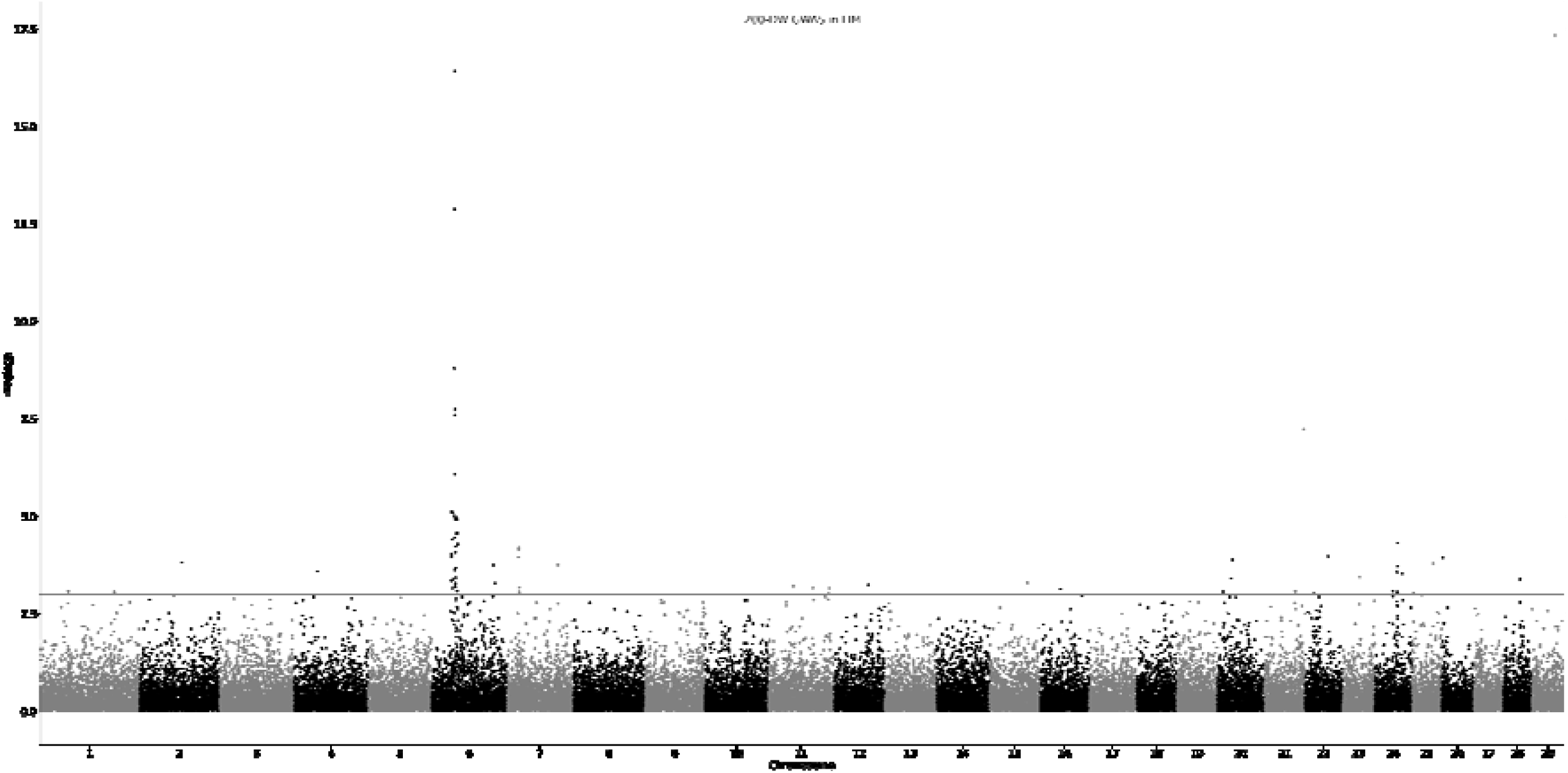
Manhattan plot (p-values) from GWAS for LIM. **Legend:** 200-DW: 200-day weight, LIM: Limousin.

For the HOL evaluation with a larger reference size, the SNP reliability in the HOL evaluation ranged between 0.01 and 0.92, with an average of 0.42. There is a clear difference between the average SNP reliabilities and the LIM and HOL evaluations. This can be attributed to the size of the reference population. In the LIM evaluation, non-zero SNP reliabilities ranged between 0.005 and 0.05, with a mean of 0.02. Figure 2 shows the Manhattan plot for the SNP effects obtained from the single-step evaluation of production traits in the HOL evaluation. Figure 3 shows the Manhattan plot of p-values for the four traits in the HOL evaluation. Similar to the LIM evaluation, we used an arbitrary threshold of 0.10% of total genetic variance explained by a single marker [9]. Using the method described in this study, we detected 2239 SNPs across all chromosomes with a statistically significant (p < 0.05) influence on MY. Using the Bonferroni adjusted threshold of significance (p < 0.05/45613), 106 SNPs on 22 chromosomes had a statistically significant effect on MY. Of these 106 SNPs detected using the p-value approach, 42 SNPs across 13 chromosomes were identified using the percentage of variance explained approach. 24 significant SNPs were on BTA 14. 16 of these SNPs (BTA 14:1463676 - 2909929) were detected around the DGAT1 region [30]. A second set of SNPs were identified on BTA 14: 66118162 – 68225066. Table 2 shows 19 SNPs on 10 chromosomes that were identified to significantly influence MY, FY and PY. 5 of these SNPs on BTA 2:128670186, 11:665543, 12:56367819, 18:577447 and 23:34516184 also had significant effects on SCS.

**Table 2:**
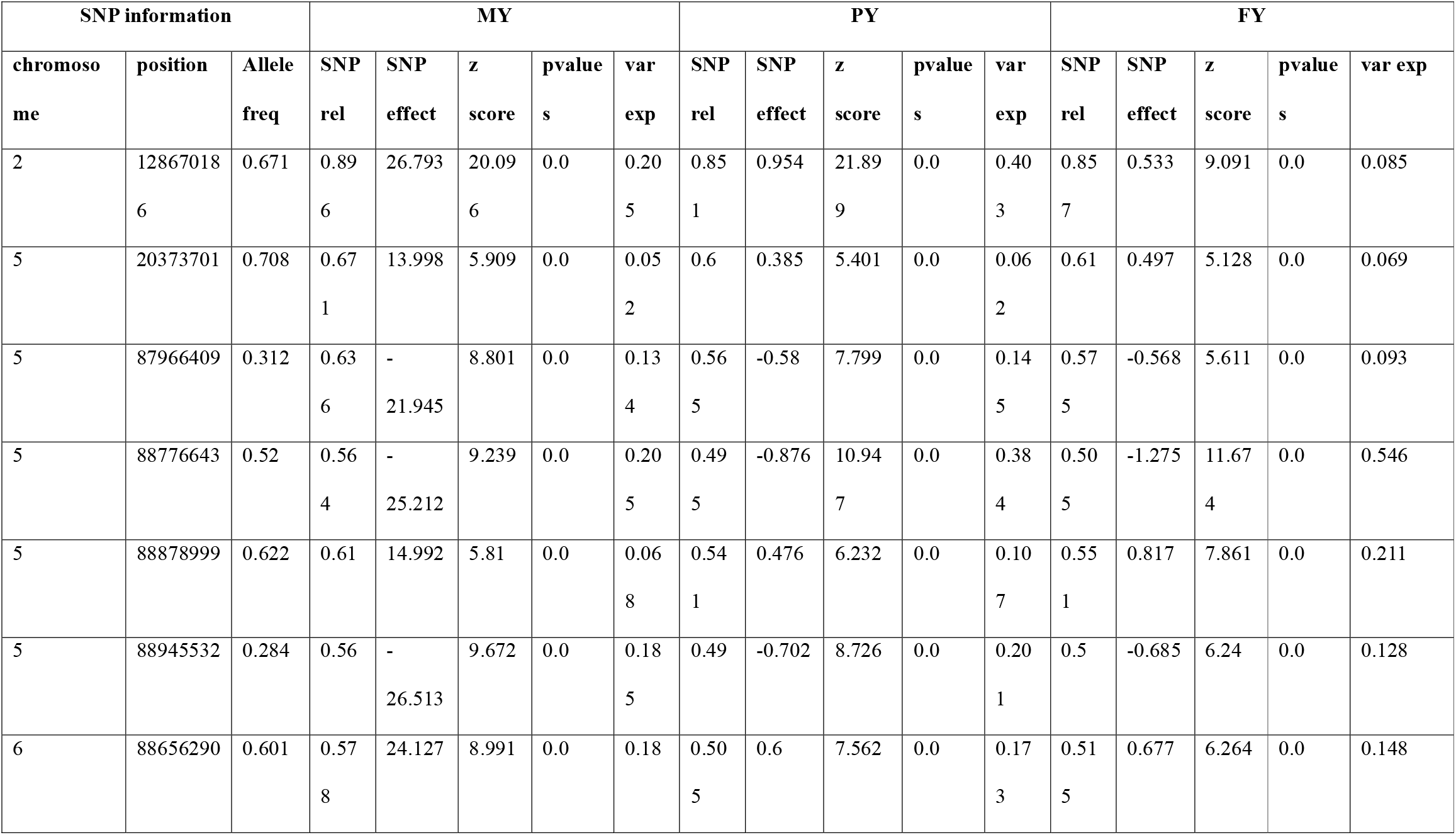
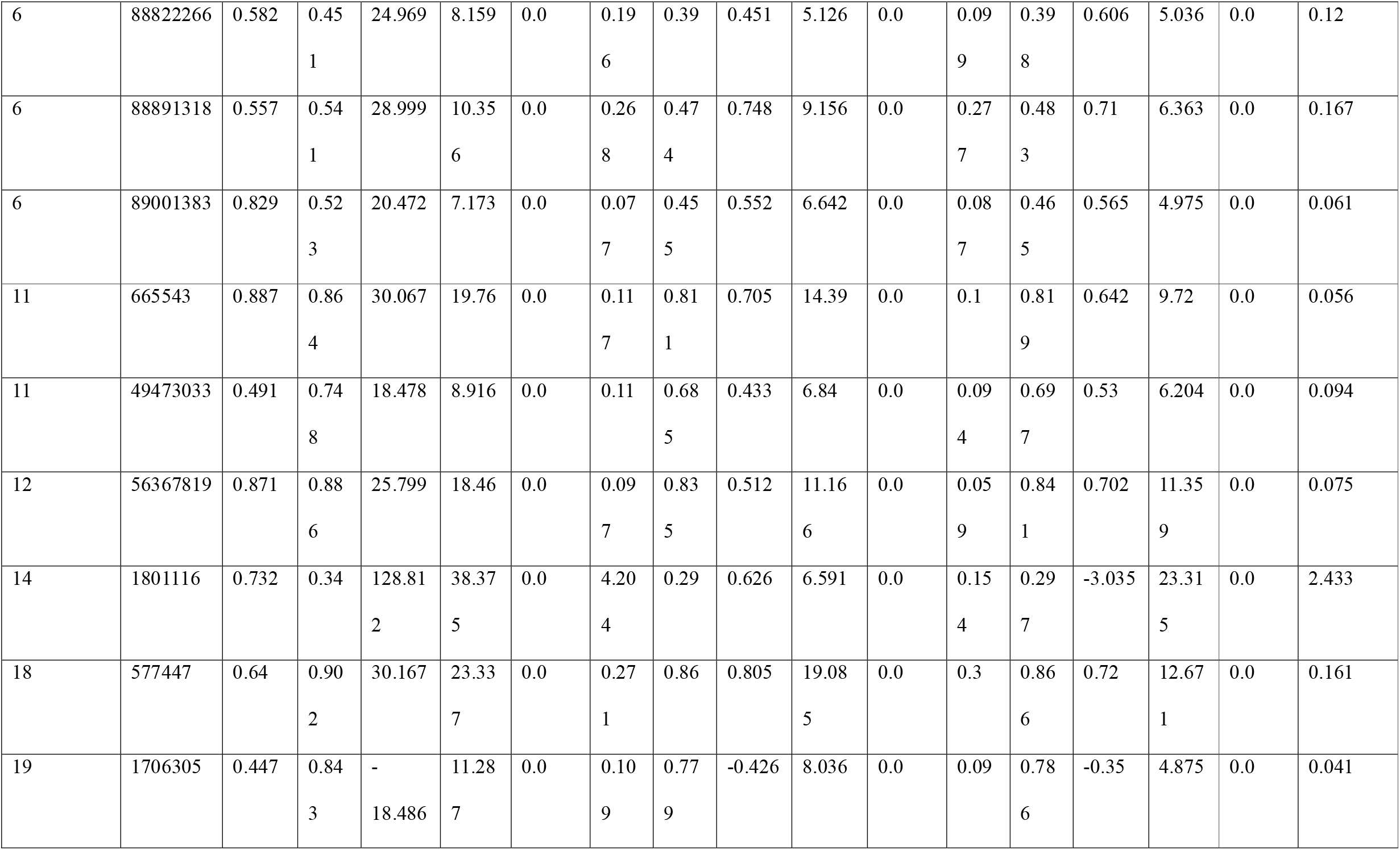
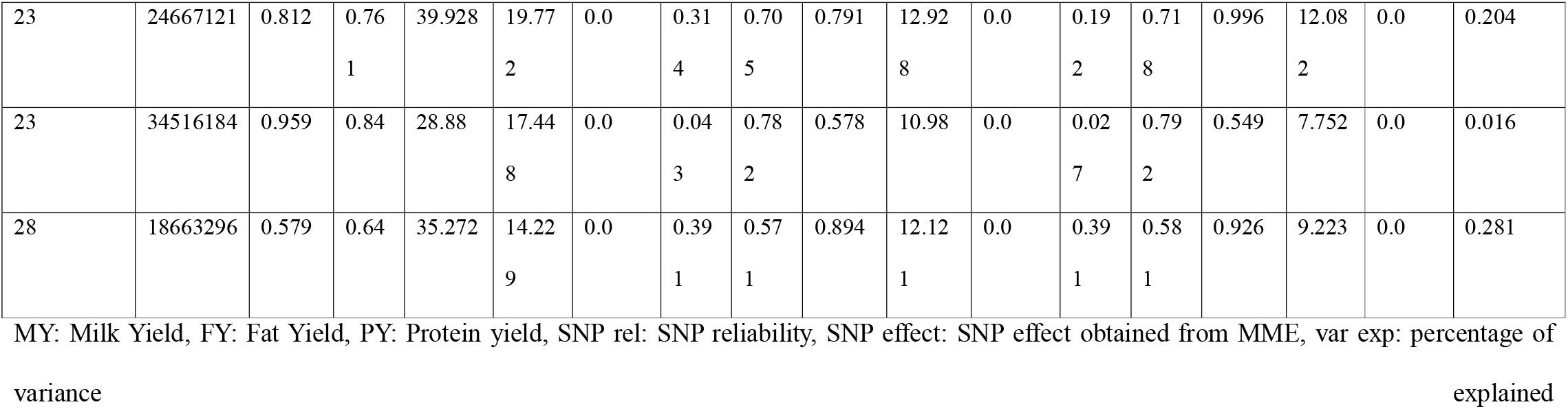
Significant SNPs on in MY, PY, FY.

For the traits MY, PY, and FY, all SNPs identified in the percentage of variance approach were also identified using the Bonferroni adjusted threshold. For SCS, three SNPs on chromosome 6:88,592,295 – 88,891,318 and one on chr 18:44,524,702 were identified only in the percentage variance-explained approach. For the SNP on chromosome 18, another SNP in the region 18:577,447 was detected in the p-value approach. Although the three SNPs identified on chromosome 6 in the percentage of variance explained approach explained between 0.10 and 0.13 percent of genetic variance, they were not significant with the Bonferroni adjusted threshold of significance. However, with a lower threshold of 0.05, these SNPs were significant. Many of the regions identified in the HOL evaluation have been identified in conventional GWAS of production traits across several dairy cattle populations [30–32].

Figure 5 and Fig. 6 show the distribution of the SNP reliabilities across the chromosomes for 200-DW in LIM and MY in HOL, respectively. The SNP reliability from the LIM evaluation ranged between 0.01 and 0.03, reflecting the reference population’s small size. The HOL evaluation with a considerably larger reference population has an average SNP reliability ranging between 0.32 in SCS and 0.42 in MY.

**Figure 5:**
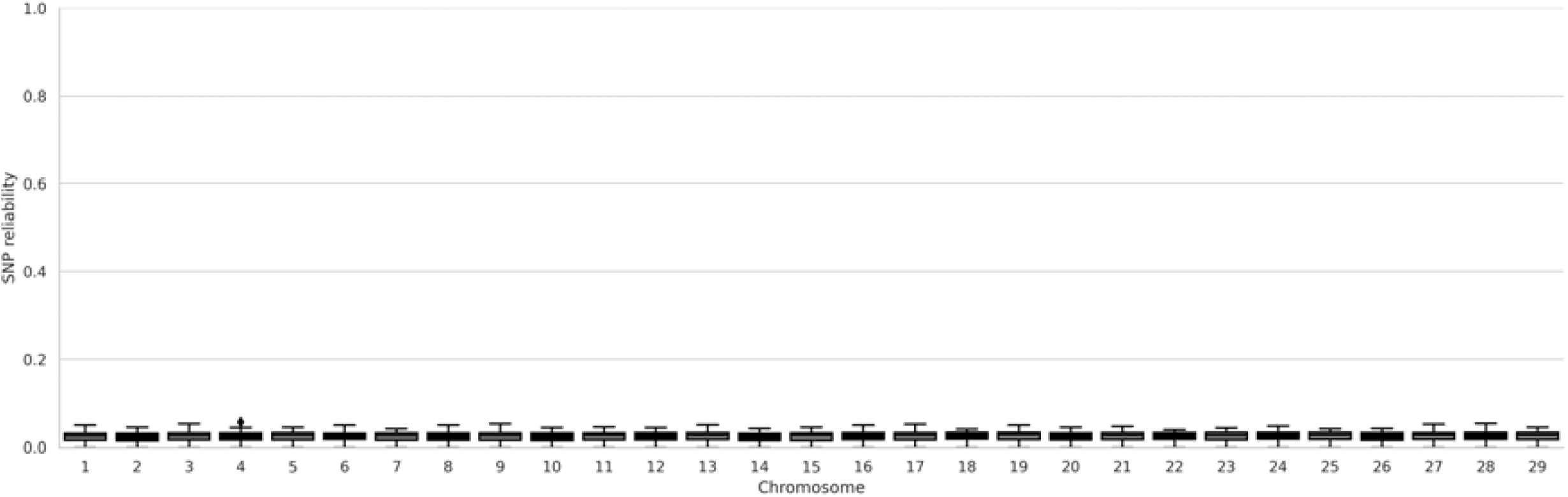
Distribution of SNP reliability by chromosome in LIM.

**Figure 6:**
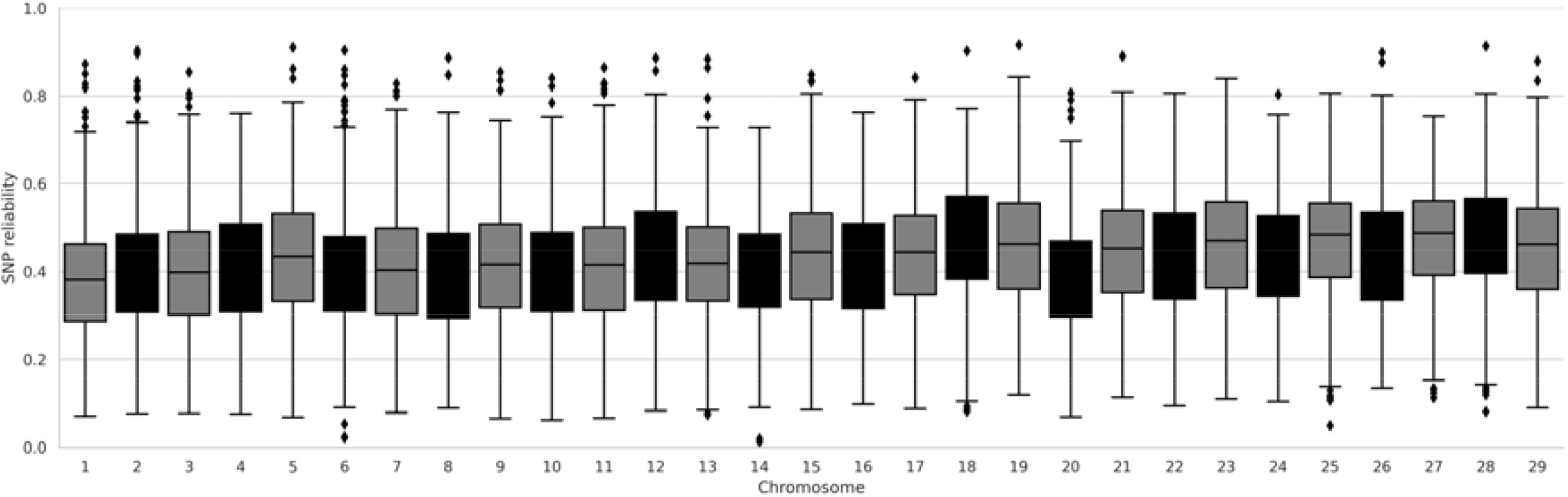
Distribution of SNP reliability by chromosome (MY)

## Discussion

Single-step methods simultaneously evaluate genotyped and ungenotyped animals in a single genomic analysis. Single-step implementations have routinely published the SNP effects estimates. However, due to the absence of methods to compute p-values of SNP effects in single step approaches, many studies have not included statistical tests to identify significant SNPs. Aguilar [9], introduced an approach to define p-values in the ssGBLUP model. However, to our knowledge, there has been no approach for the ssSNPBLUP framework nor a systematic evaluation of the p-values in large and small reference populations. This study introduces an approach to calculate p-values for the ssSNPBLUP framework. The regions identified by the method show good empirical agreement with known associations reported in studies on dairy and beef cattle in the literature. The results from this study indicate a severe underestimation of the p-values estimated in the small reference population. This was attributed to the very low SNP reliability obtained from the LIM population with small reference population size. The method introduced in this study was compared to a conventional single marker regression GWAS study. The marker effects obtained from the ssSNPBLUP and single marker regression approach both signified a region of interest on chromosome 6. However, due to the low number of individuals in the reference population, this region on chromosome 6 was not significant using the approach introduced in this study. The result from this study indicate that the p-value obtained from ssSNPBLUP models may by significantly underestimated or the p-values obtained from the GWAS model are overestimated.

Comparing the SNPs identified as significant in the GWAS, variance explained and ssSNPBLUP approaches demonstrate that the p-values obtained using the ssSNPBLUP framework may be severely underestimated in populations with small reference sizes, particularly when SNP reliability observed in these populations is low. While both methods identified a region of interest on chromosome 6 associated with the 200-DW in the LIM evaluation, the ssSNPBLUP-derived p-values did not reach predefined threshold of significance. Conversely, the GWAS approach identified SNPs in this region as significant (see Fig. 4).

In the LIM evaluation, SNP reliabilities ranged between 0.005 and 0.05, with an average of 0.02. In contrast, the HOL evaluation, which utilized a larger reference population, exhibited significantly higher SNP reliabilities, ranging from 0.01 to 0.92, with a mean of 0.42 in MY. The disparity in SNP reliabilities underscores the critical role of reference population size in determining the accuracy, certainty and reliability of the SNP effects obtained in genomic evaluations. This study introduced a novel method to calculate p-values within the ssSNPBLUP framework and systematically compared it to a conventional single-marker regression GWAS approach using the LIM dataset. Unfortunately, due to the large size of the reference population, a comparative single marker regression GWAS could not be carried out in the HOL evaluation. Further research should be undertaken to study the effects and interactions of reference size, allele frequencies and QTL effects on the ability of this method to detect significant SNPs.

An important outcome of this study is the empirical agreement between regions identified using the ssSNPBLUP framework, conventional single marker regression GWAS and previously reported QTL in dairy and beef cattle populations. In beef cattle, several studies [26–29] have reported QTLs on chromosome 6 affecting body weight, growth, and carcass traits. This study identified SNPs in the same region using ssSNPBLUP approach, although their statistical significance was lower than the predefined threshold of significance. In the HOL evaluation, the use of a larger reference population facilitated the detection of significant SNPs across various production traits, including MY, PY, and FY. Notably, the percentage of variance explained approach identified SNPs that were also confirmed by the Bonferroni-adjusted threshold. Consistent with findings from prior GWAS studies on production traits in dairy cattle, regions such as DGAT1 on chromosome 14 were highlighted. For SCS, significant SNPs were identified using the percentage of variance approach and the less stringent threshold of 0.05. However, these SNPs were not significant in the more stringent Bonferroni adjusted threshold. At a significance threshold of p < 0.05, a large number of SNPs were identified as significant in the HOL evaluation, (MY: 2,239, PY: 1,629, FY: 964, SCS: 425). A more stringent threshold (p < 0.05/45613) resulted in (MY: 106, PY: 39, FY: 42, SCS: 8) significant SNPs. The differences between the number of significant SNPs highlight the need for flexible thresholds that will depend on the aim of the study carried out. The specified threshold parameter should also consider population-specific factors such as reference size and trait heritability and number of markers. Considering the infinitesimal allele model, SNP markers would explain average SNP variance *average* = *var*(*DGV*)/ Σ 2*pq*. With this in consideration, the definition of what constitutes as a significant SNP can only be defined based on the aim of the study. Going forward, studies may choose to consider the definition of significance based on the presence of a major QTL in linkage disequilibrium with the marker or the estimated effect of a SNP marker being different from zero.

The study also emphasized the utility of SNP reliability as an indicator of accuracy of SNP effects obtained from single step genomic evaluations. The stark contrast in SNP reliability between the LIM and HOL populations highlights the challenges of conducting robust genomic analyses in populations with limited reference data. The low SNP reliabilities in the LIM population likely contributed to the observed underestimation of p-values, underscoring the importance of increasing reference population size to improve the precision of genomic evaluations and particularly reducing the statistical uncertainty associated with the estimation of SNP effects.

In comparing SNP effects in genetic standard deviations between the HOL and LIM evaluations, the larger reference size in HOL enabled more precise estimation of SNP effects and higher reliability. This aligns with the general understanding that larger reference populations enhance the statistical power to detect true genetic associations. Computationally, the ssSNPBLUP approach, in which computational cost is limited by the number of markers also offers advantages in the estimation of SNP effects over the ssGBLUP and conventional GWAS approaches which depend on the number of genotyped individuals. Although, approaches such as Algorithm for Proven and Young (APY) [33,34] have been introduced to manage the computational demands in the ssGBLUP framework, the ssSNPBLUP framework provides advantages in estimating SNP effects and their associated uncertainties using all information from large datasets such as those available in many national databases. Further investigations should address comparing the SNP effect estimates and their reliabilities derived from back solving GEBVs to SNP effects obtained in the ssGBLUP-APY approach to the SNP effects obtained from the ssSNPBLUP framework. Evaluating the impact of the number and collection of core animals in the APY approach on SNP effects and their significance will also be important aspect to study.

In this study, the mixed model Eq. (6) does not take RPG into account when calculating reliability values of the SNP effect estimates via Eq. (8), although the RPG is considered afterwards when computing standard errors of the SNP effect estimates (Eqs. (10) and (11)). Ideally, the RPG could be included in the mixed model Eq. (6) as shown in [19,35]. However, the computation would have become too heavy and infeasible, due to the number of reference animals in the HOL dataset. Future research should be focused on developing more efficient statistical methods to estimate reliability values of the SNP marker effect estimates by including the RPG effect directly into the mixed model Eq. (6).

### Implications and Future Directions

This study provides critical insights into the application of ssSNPBLUP for genomic evaluations across populations with varying reference sizes. The underestimation of p-values in small populations necessitates further research to refine the methodology and improve its robustness. In this study, we have used real data to test our approach to detecting SNP markers that significantly influence traits in dairy and beef cattle datasets. We attempted to validate many of the SNPs we identified against previously published markers identified using conventional single marker GWAS approaches. However, future studies should focus on controlled simulation studies to further understand the effect and interaction of reference size, allele frequencies and the segregating QTL effects on the ability of GWAS and single step frameworks models to detect genomic regions significantly influencing the trait.

## Conclusions

In conclusion, the ssSNPBLUP framework offers a powerful tool for integrated genomic evaluation, but its application requires careful consideration of reference population size, SNP reliability, and statistical thresholds. By addressing these challenges, the utility and accuracy of ssSNPBLUP can be further enhanced, benefiting genomic evaluations in both large and small reference populations.

## Declarations

### Ethics approval and consent to participate

Not applicable

### Consent for publication

Not applicable

### Availability of data and materials

The data supporting this study’s findings may be available upon request from vit Verden. However, restrictions apply to the availability of these data, which were used under a licence of a material transfer agreement for the current study and thus are not publicly available.

## Authors’ contributions

DA, ZL, DS, and JT conceived the study. DA and ZL wrote the code for the analysis. DA, HA, and ZL, analysed the result of study. ZL, GT and JT coordinated the study. All authors contributed to the manuscript’s writing. All authors read and approved the final manuscript.

## Acknowledgements

This study received no external funding. The authors have not stated any conflicts of interest.

